# Chop-n-drop: *in silico* assessment of a novel single-molecule protein fingerprinting method employing fragmentation and nanopore detection

**DOI:** 10.1101/2021.05.27.445917

**Authors:** Carlos de Lannoy, Giovanni Maglia, Dick de Ridder

**Affiliations:** Bioinformatics Group, Wageningen University, 6708PB, Wageningen, The Netherlands; Groningen Biomolecular Sciences Biotechnology Institute, University of Groningen, 9747AG Groningen, The Netherlands

## Abstract

The identification of proteins at the single-molecule level would open exciting new venues in biological research and disease diagnostics. Previously we proposed a nanopore-based method for protein identification called chop-n-drop fingerprinting, in which the fragmentation pattern induced and measured by a proteasome-nanopore construct is used to identify single proteins. However whether such fragmentation patterns are sufficiently characteristic of proteins to identify them in complex samples remained unclear. In the simulation study presented here, we show that 97.9% of human proteome constituents are uniquely identified under close to ideal measuring circumstances, using a simple alignment-based classification method. We show that our method is robust against experimental error, as 78.8% can still be identified if the resolution is twice as low as currently attainable and 10% of proteasome restriction sites and protein fragments are randomly ignored. Based on these results and our experimental proof-of-concept, we argue that chop-n-drop fingerprinting has the potential to make cost-effective single-molecule protein identification feasible in the near future.

## 1 Introduction

Over the past decades, mass spectrometry (MS) has allowed for ground-breaking discoveries in proteomics, enabling such impressive feats as the definition of a human protein atlas [1] and large-scale screening for protein disease biomarkers [2]. However, not all protein-related research questions may be addressed by MS. Examples are found in the nascent field of single-cell proteomics which, following the example of single-cell transcriptomics, is expected to give unprecedented insight into cell functioning and pathology [3]. While MS has already made strides in this field by enabling the detection of proteins present at thousands of copies per cell [4], some important and clinically relevant proteins such as signaling molecules and transcription factors are expected to be present in the range of dozens of copies [5]. The development of novel single-molecule protein identification methods is therefore necessary to unlock the true potential of single-cell proteomics.

In the search for single-molecule alternatives to MS, two main venues are currently being explored. On the one hand, conceptual methods utilizing the read-out of fluorescent dyes attached to a subset of residue types have shown promising results [6, 7, 8]. However, methods using fluorescence-based readout strategies require efficient and specific labeling of residues. Optimizing labeling strategies is non-trivial [e.g. 9, 10] and less-than-perfect labeling may decrease accuracy, thus a label-free method would be preferred. On the other hand, electrical readout of protein properties, either folded or unfolded, may be generated by feeding the protein through a nanopore, over which an electrical potential is applied [11, 12]. Similar to nanopore sequencing of DNA, changes in the pore’s electrical resistance while a protein is passing may give information on its properties.

In prior work, we showed that engineered complexes of FraC nanopores and proteasomes can be readily assembled without loss of proteasome activity or electrical conductance of the pore [13]. Furthermore, we have shown that a linear relation exists between residual current through FraC pores and the molecular weight of passing protein fragments [14]. We thus proposed that proteasome-nanopore constructs can be used to identify proteins, in a conceptual method dubbed chop-n-drop fingerprinting [13]. An unknown protein can be processed terminal-to-terminal by the construct, cleaving it at proteasome target sites, after which the molecular weight of sequentially released fragments can be estimated based on the residual electrical current as they pass through the nanopore. The sequence of measured fragment weights can then serve as a characteristic signature – a fingerprint – of the protein. Once proven, this fingerprinting method can easily be implemented in a highly parallel fashion by adapting existing hardware that was developed for nucleic acid sequencing. Compared to both MS and existing fluorescence-based measurement equipment, this hardware is inexpensive and has a small benchtop footprint, thus opening up opportunities for field diagnosis and in-house analysis for even small laboratories. It is as of yet however unclear whether chop-n-drop fingerprints are sufficiently characteristic to identify a single protein in highly complex mixtures.

Here we present a computational analysis of the chop-n-drop method, in which we show that simulated fingerprints of all proteins in the UniProt human proteome can be accurately classified using a simple alignment-based method. Considering these and previously published experimental results, we argue that chop-n-drop fingerprinting is a promising concept for cost-effective single-molecule protein identification.

**Figure 1:**
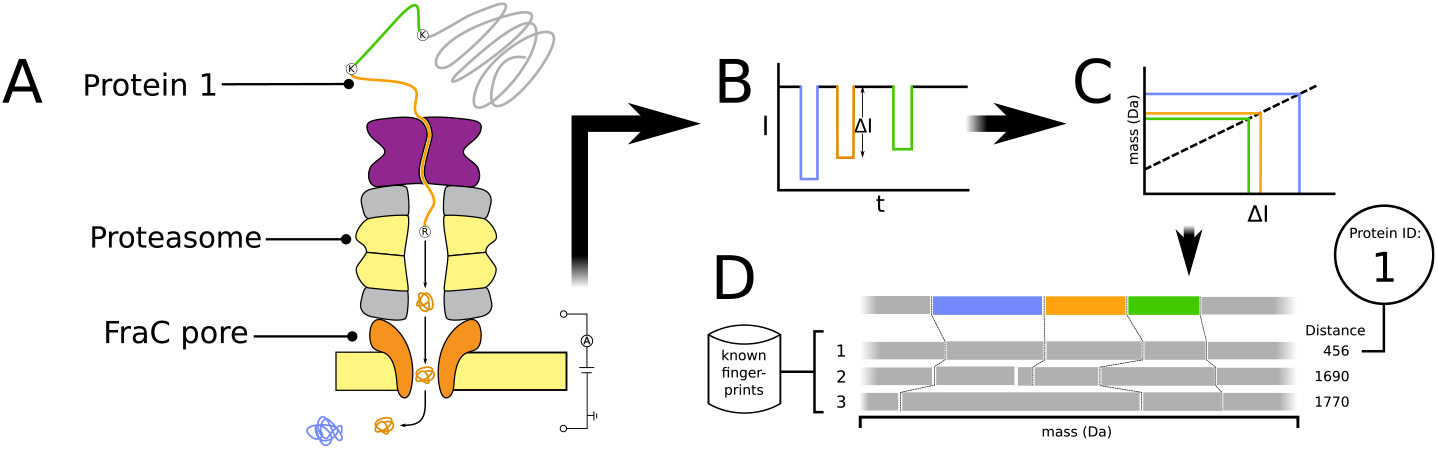
Schematic overview of the chop-n-drop fingerprinting method. **(A)** A protein is fragmented by a proteasome directly introduced above a nanopore. The protease is engineered to lyse proteins at particular residues. **(B)** As the fragments pass the pore, a change in electrical current through the pore is measured. **(C)** The molecular weights of the fragments are estimated from the magnitudes of the current changes. **(D)** Finally the produced sequence of fragment weights is aligned to database fingerprints of known proteins, to identify the protein.

## 2 Results

### 2.1 Simulation and classification method

To estimate the performance of the chop-n-drop fingerprinting method on a highly complex protein identification task, we developed a simulation pipeline mimicking the experimental procedure, including several sources of biological and technical noise that we expect to encounter.

In essence, the chop-n-drop fingerprint of a protein only consists of a sequence of weights, which are deduced from pore current blockades caused by sequentially cleaved-off fragments passing through the nanopore. The simulation of this process follows a straight-forward two-step process. First, akin to the proteasome cleaving a protein into fragments, we divide a given protein sequence into sub-sequences by splitting it at the proteasome’s target sites. Although engineering proteasome specificity in our system is still a work in progress we assume here that we can force it to exhibit only trypsin-like behavior, thus we split sequences after arginines and lysines, unless followed by a proline. To account for the fact that the proteasome will likely fail to cleave at a fraction of target sites, we only cleave each target site with a certain probability, which we refer to as the proteasome efficiency (*e_p_*).

Subsequently we mimic the passing of fragments through a heptameric FraC pore, the readout of the current blockade and the estimation of the fragment weight, by simply translating the sub-sequences into corresponding fragment weights. Although weights can be determined with high precision from sequences, the measurements in experiments may be less accurate and marked by a given resolution (*r*), the smallest detectable weight difference. In experimental setups, this parameter is dependent on pore and measuring equipment properties. To account for this in simulations, Gaussian noise is added to fragment weights, where the standard deviation of the noise is related to *r* (see Methods). Fragments weighing less than 500Da are removed, as they typically escape detection of heptameric FraC nanopores [14]. Furthermore, as we previously found that the relation between weights above 2kDa and current blockades is non-linear[14], all fragment weights larger than this value are reduced to 2kDa. Lastly, although we expect the seal between proteasome and pore to be extremely tight based on molecular dynamics simulations [13], fragments may fail to enter the pore after cleavage. We account for this by only retaining each fragment with a certain probability, which we refer to as the capture rate (*C*). Although *C* is likely dependent on the size and charge of individual fragments, the relationship between these factors is unclear, thus we assume *C* to be constant. The resulting sequence of fragment weights returned by this process constitutes the fingerprint for a protein.

We used fingerprints generated using our pipeline to develop a classification method, which assigns a protein identity to a given fingerprint. We follow an alignment-based approach, where a query fingerprint is aligned to a database of previously generated fingerprints, using a custom dynamic programming implementation (Supplementary figure S1, see Methods). The database fingerprint that is most similar to the query fingerprint is assumed to have come from the same protein.

### 2.2 Simulations under optimal conditions

We ran our simulation pipeline and classification method on all sequences in the UniProt human proteome (*n* = 20, 395). Under close to ideal simulated noise parameters (*e_p_* = 0.99, *r* = 5.0Da, *C* = 0.99) we find that our alignment based approach retrieves the correct identity for 97.9% of fingerprints (Figure 2). Inspection of made alignments shows that our algorithm correctly handles missing and fused fragments (Supplementary figure S2A). The majority of misclassifications occurs for shorter proteins, under 250 residues in length. Of misclassified fingerprints, 48% shows more than 80% amino acid sequence identity to the protein as which it was wrongly identified, indicating that the resolution of 5Da assumed here is insufficient to consistently separate such similar entities (Supplementary figure S3). Upon inspection of these cases, we find that many misclassifications were in fact mix-ups between paralogous sequences. The remaining misclassifications are caused by chance alignments with different fingerprints (Supplementary figure S2B). This is expected to occur more often if a protein is shorter, as it will generally produce a fingerprint of fewer elements, which is less likely to yield a unique pattern.

**Figure 2:**
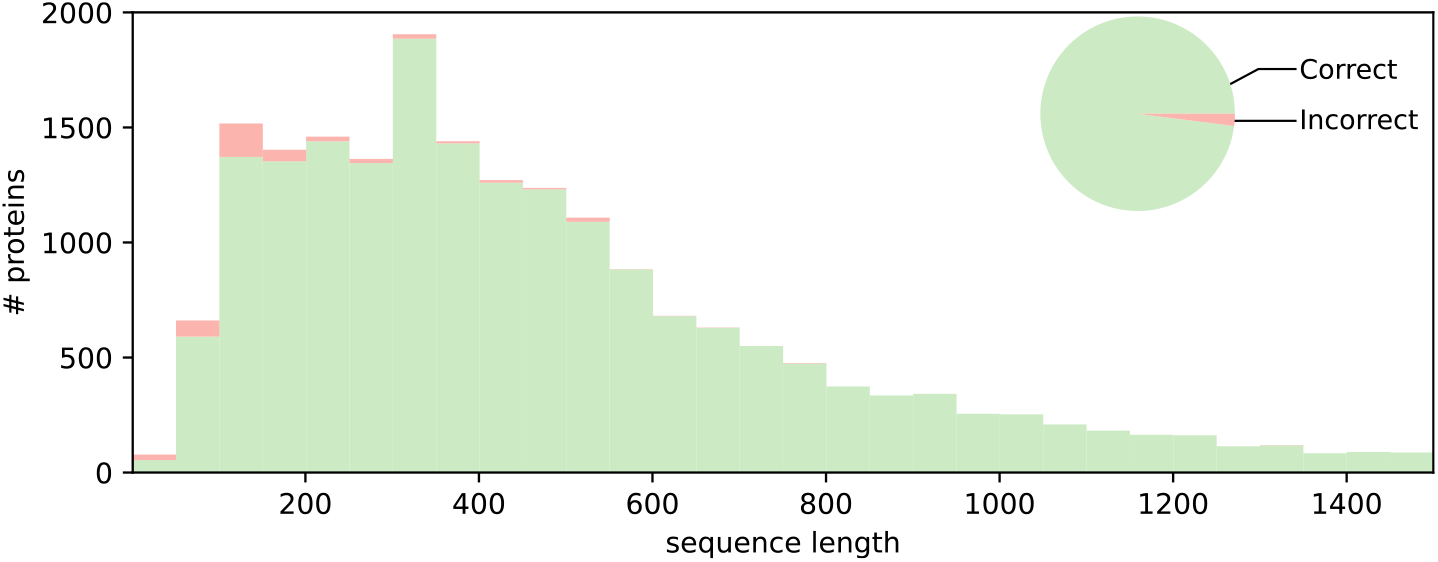
Cumulative histogram of correct and incorrect classifications of simulated chop-n-drop protein fingerprints for all human proteome constituents, assuming low noise parameters; resolution *r* = 5Da, capture rate *C* = 0.99 and proteasome cleaving efficiency *e_p_* = 0.99. Numbers are shown distributed over sequence length (bars), and relative to the total number of proteins (pie chart).

### 2.3 Simulations under suboptimal conditions

We subsequently probed how resistant chop-n-drop fingerprinting is to higher levels of experimental noise, by varying one noise parameter at a time while keeping all others near their optimal values (*e_p_* = 0.99, *r* = 5.0Da, *C* = 0.99). In each case we find that accuracy deteriorates gracefully with parameter value (Figure 3A). Interestingly, we still attain an accuracy of 92.6% at a resolution of 50Da, which is worse than the 40Da resolution we reported previously [14] and more than tenfold worse than our current-best resolution of 4Da (GM, unpublished results). Similarly, we find that a lower proteasome efficiency or catch rate of 90% still results in 93.6% and 90.7% accuracy on average respectively.

**Figure 3:**
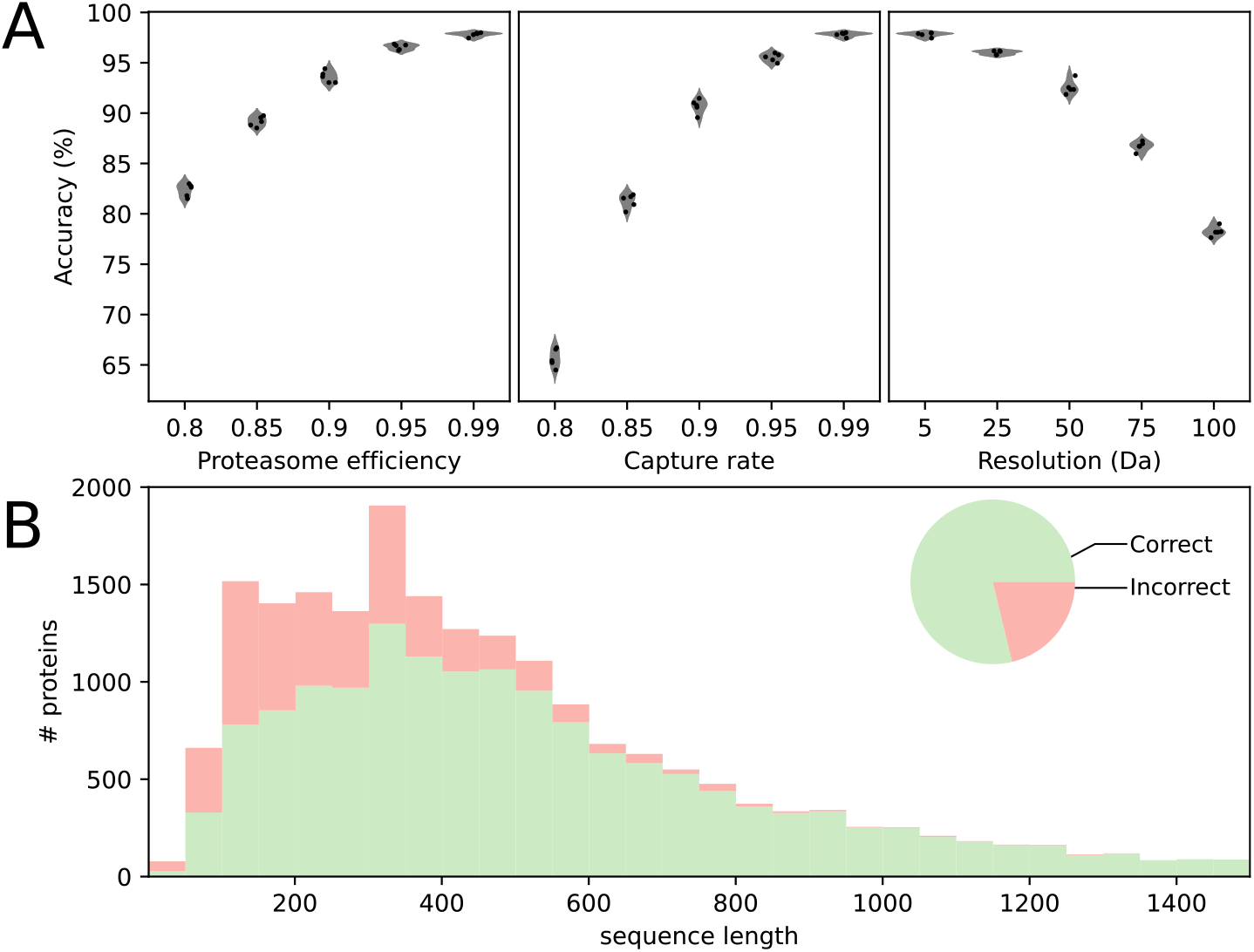
**(A)** Fingerprint classification accuracy over a range of sub-optimal noise parameter values; resolution (left), capture rate (mid) and proteasome efficiency (right). For each case the unvaried noise parameters are set to near-optimal values (capture rate *C* = 0.99, resolution *r* = 5.0Da and proteasome efficiency *e_p_* = 0.99). Five replicates were generated for each parameter combination. **(B)** Cumulative histogram of correct and incorrect classifications of simulated chop-n-drop protein fingerprints for all human proteome constituents, assuming more realistic noise parameters; *r* = 10Da, *C* = 0.90 and *p_r_* = 0.90. Numbers are shown distributed over sequence length (bars), and relative to the total number of proteins (pie chart).

Finally, we repeated a simulation on the entire dataset with all noise parameters at sub-optimal values (*e_p_* = 0.90, *r* = 10.0Da, *C* = 0.90). Even under these circumstances, we find that 78.8% of proteins are correctly classified (Figure 3B). Here too, it should be noted that most incorrectly classified proteins were of lower sequence length.

## 3 Discussion

Single-molecule (SM) protein fingerprinting holds great promise to revolutionize biological research and diagnostics [15]. We have previously proposed that this may be accomplished using a novel proteasome-nanopore construct, which cleaves a target protein into fragments and subsequently reads out the fragment weights [13]. Here we present simulation results indicating that the produced sequence of fragment weights contains sufficient information to identify a protein.

In the presented simulations, we included sources of noise that may hamper fingerprint measurements in practice. We assumed that the proteasome may not cleave each target site, that weight measurements may be inaccurate up to a given weight resolution and that not all cleaved-off fragments may be caught in the nanopore. Assuming higher noise parameter settings – a fragment capture rate and proteome efficiency of 90%, with a measurement resolution of 10Da – for each of these noise sources, we find that overall accuracy remains sufficiently high at 78.8%. As accuracy increases with protein length, we find that chop-n-drop fingerprinting should be particularly suitable to identify larger proteins.

Our simulation builds on the assumption that fragment weight is correlated to the residual current measured while the fragment passes the nanopore. Indeed, we have previously shown that this is the case for selected fragments weighing between 500 and 2000Da [14]. However it should be noted that rather than the peptide’s weight, its volume, charge and shape influence residual current. Once the experimental methodology has been further developed and protein fingerprints can be measured more routinely, we can define the relation between these properties and the residual current in more detail to predict fingerprints in a more robust manner.

The existence of different proteoforms, which was not accounted for in this simulation, presents both an opportunity and a challenge to chop-n-drop fingerprinting. Through alternative splicing and post-translational modification (PTM), multiple proteoforms with different functions may be generated from the same gene [16]. Depending on the spliceoform or the PTM types present, different proteoforms may generate distinct fingerprints. This allows their individual identification at SM resolution, which is an important potential application of SM analysis, but also adds tens of thousands of potential fingerprint patterns, which further complicates the task of fingerprint classification. A solution may be to fractionate samples prior to chop-n-drop analysis, after which each fraction may be analysed using a dedicated classifier which only considers the proteoforms that could be present in a given fraction.

Over the past years, the obstacles on the road toward SM protein fingerprinting have been attacked vigorously from multiple angles, with several groups showing promising initial results and proofs-of-concept. While each proposed method has shown particular strengths, we argue that chop-n-drop combines several properties not found together in other methods. First, unlike fluorescence-based methods [7, 6, 8] it does not require the implementation of any labeling chemistries as properties of the target protein are read out directly, thus evading issues with erroneous labeling and simplifying sample preparation. As a trade-off, fluorescence-based methods are more sensitive to differences between proteoforms as long as the difference involves the position or presence of a targeted residue type. As we show here that even at high resolution our method misclassifies proteins with high sequence similarity to other entries, it is likely that differences between highly similar proteoforms may also remain unnoticed.

Different methods based on the readout of folded proteins by electrical current blockage of a nanopore have been proposed as well [12, 17, 18]. These were unable to analyse a wide range of protein sizes however; as the pore lumen needs to be of an appropriate volume for the analysis of a given protein size, a single nanopore is not able to detect minute differences in both small and large proteins. Here this problem is mitigated by the fragmentation step.

Most importantly however, the hardware required to implement chop-n-drop fingerprinting in a highly parallelized setting can be readily borrowed from commercial platforms for DNA sequencing using nanopores, which are inexpensive and have already been miniaturized to a handheld format. As such we envision that our method could soon fill a niche that no other method currently can; that of small-scale, in-house single-molecule protein identification.

In conclusion, we provide evidence that chop-n-drop fingerprints can provide sufficient information to identify proteins in complex samples, and present a suitable alignment-based classification method. Upon optimization of the fingerprinting procedure, we envision that our method may see practical implementation in the near future.

## 4 Methods

Code for *in silico* fingerprint generation and classification was wrtten in Python 3.8 (Python Software Foundation, www.python.org), and is freely available at https://github.com/cvdelannoy/chop_n_drop_simulation.

### 4.1 *In silico* fingerprint generation

We generate *in silico* chop-n-drop fingerprints by splitting protein sequences at protease target sites and calculating the weights of the resulting fragments from their sequences. We assume that fragments of a weight lower than 500Da are undetectable, thus these fragments are removed from fingerprints. Fragments of a weight larger than 2kDa are set to 2kDa, as prior investigations showed that the relationship between weight and current blockage is non-linear above this weight[14].

Three parameters are set to represent different noise sources; catch rate *C*, proteasome efficiency *e_p_* and resolution *r*. The catch rate denotes the fraction of fragments that enters the pore after lysis and is measured. In our simulations each fragment is retained with a probability of *C*. The proteasome efficiency denotes the fraction of target sites at which the proteasome cleaves. In simulations, each target site has a probability of *e_p_* of being cleaved. Note that a failure to cleave will result in two fragments being fused together, after which they remain represented in the fingerprint as the sum of their weights. Finally, the resolution denotes the minimum difference in fragment weight that can still be detected by current blockage, expressed in Da. In our simulations, the resolution is represented by the magnitude of Gaussian noise added to fingerprint weights. Specifically, we define the standard deviation of the noise such that the probability of a fragment size measurement deviating *r* from its actual size is fifty percent:

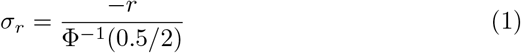

Here *σ_r_* is the standard deviation of the Gaussian noise, *r* is the resolution, and Φ^-1^ is the inverse cumulative distribution function of the standard normal distribution.

### 4.2 Simulation

We ran *in silico* digestions on all sequences in the UniProt human proteome (UP000005640). To compile a database of fingerprints with known identity, we first performed an in *silico* digestion under noiseless circumstances (i.e. *C* =1.0, *e_p_* = 1.0 and *r* = 0.0Da). Then we ran several subsequent digestions for a range of values for *C, e_p_* and *r*. Fingerprints from these runs were classified by aligning them to database fingerprints obtained from noiseless digestions.

### 4.3 Fingerprint alignment and classification

We gauge the similarity of query and database fingerprints by aligning them using a dynamic programming algorithm (Supplementary figure S1). The dynamic programming table is filled as follows:

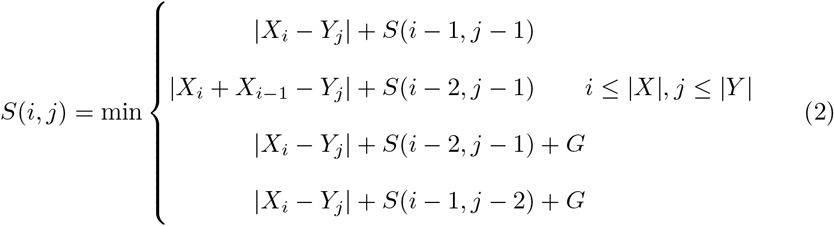

with the following conditions for edge cases to ensure that the alignment is global:

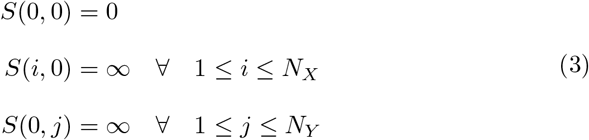

Here *S*(*i,j*) is the distance between query and database fingerprints *X* and *Y* respectively, up to fragments *X_i_* and *Y_j_* and *G* is a gap penalty. *N_X_* and *N_Y_* are the numbers of fragments in *X* and *Y* respectively. At each step in the alignment one of three actions may be taken. First, a single fragment of each fingerprint may be aligned, in which case the absolute difference of their weights is added to the total score. Second, two fragments of *X* may be aligned to one fragment of *Y*, corresponding to a missed proteasome target site. This action increases the score by the difference between the summed weight of the former and the single weight of the latter. Third, a gap may be introduced in either *X* or *Y* at the cost of a penalty. The gap penalty *G* is dependent on the resolution used during digestion:

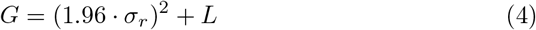

Here *σ_r_* is the resolution-dependent standard deviation of Gaussian noise added to fragment sizes during *in silico* digestion (equation 1) and *L* is the lower detection limit (*L* = 500 Da). This means that introducing a gap is preferred over matching fragments if the difference between fragment weights exceeds the difference expected in 95 percent of correct matches. The addition of *L* is required to ensure that a match is still preferred if a normally undetected fragment (i.e. of which the weight is under *L*) is fused to another fragment due to a missed proteasome target site.

A query fingerprint is classified by aligning it to all fingerprints in the database and assigning it the identity of the database fingerprint to which the distance is smallest.

## 5 Acknowledgements

This work was supported by the Vrije Programma of the Foundation for Fundamental Research on Matter under grant number 16SMPS05.

## Supplementary figures

**Figure S1:**
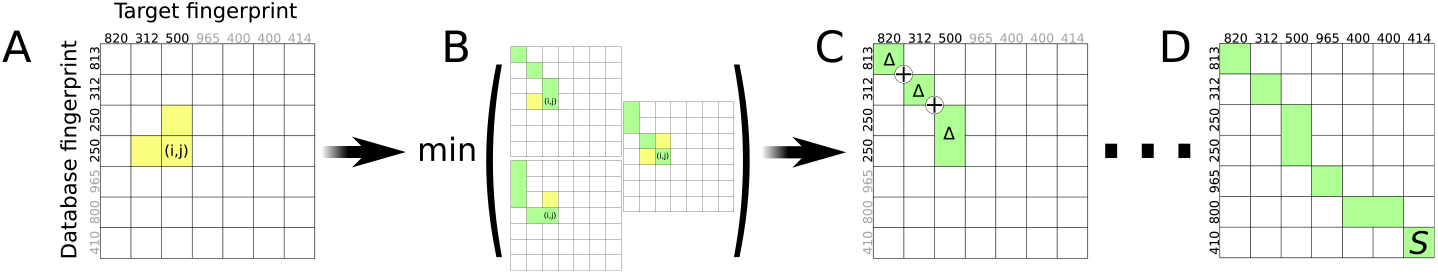
Diagram of the dynamic programming algorithm used to align database and query chop-n-drop fingerprints. **(A)** Numbers on the axes of the matrix denote fragment weights in Da. Weights involved in the displayed alignment step are black. The cells in the comparison matrix are filled row-wise starting from the top-left, by entering the alignment distance up to cell (*i,j*) in (*i,j*). **(B)** To fill square (*i,j*), two fragments of one fingerprint may be aligned to one fragment in the other, or single fragments may be aligned. A gap may also be introduced at a resolution-dependent penalty (not shown). **(C)** The option minimizing the summed distances of aligned fragments (Δ’s) up to (*i,j*) is chosen – in this case, two fragments of of the database fingerprint are aligned to a single fragment of the target fingerprint. **(D)** The process is continued until the bottom right cell is filled. The value in this cell (S) is the alignment distance for this pair of fingerprints.

**Figure S2:**
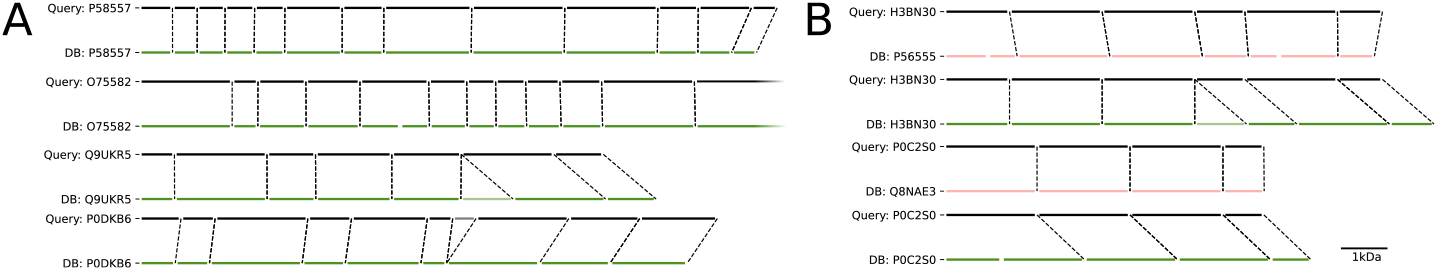
Representative examples of alignments between chop-n-drop query fingerprints and the best-matching database fingerprints. Dotted lines denote aligned fragments. Fragments for which a gap was introduced are greyed-out. **(A)** Four correct alignments. **(B)** Two incorrect alignments (red) and corresponding correct alignments (green).

**Figure S3:**
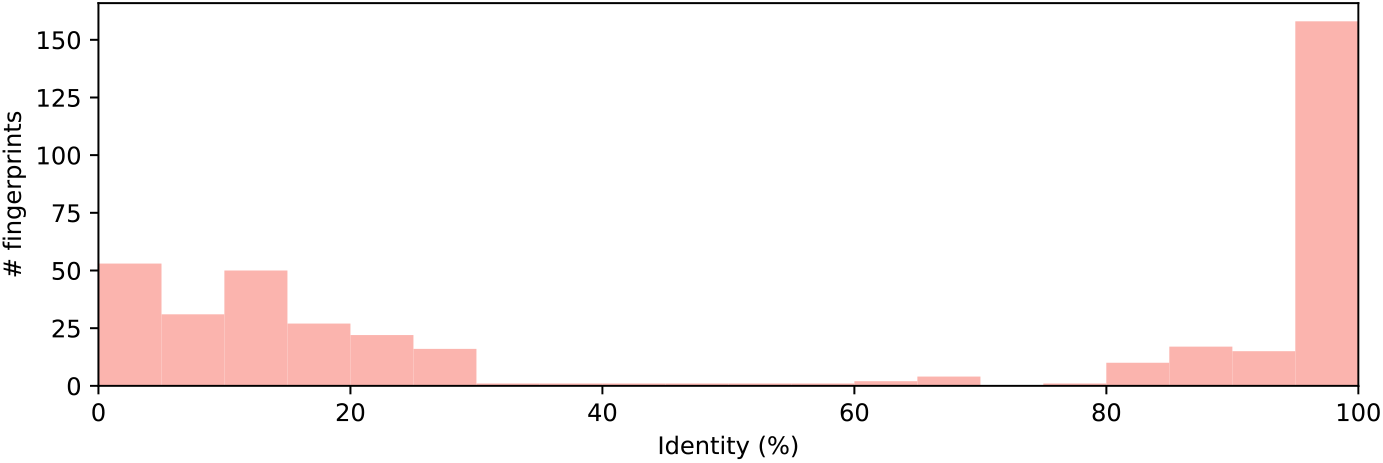
Distribution of sequence identities between misclassified proteins and the proteins for which they were mistaken based on their chop-n-drop fingerprint.

## References

[1] Fredrik Pontén, Karin Jirström, and Matthias Uhlen. The Human Protein Atlas—a tool for pathology. The Journal of Pathology, 216(4):387–393, 2008.

[2] Hasmik Keshishian, Michael W Burgess, Harrison Specht, Luke Wallace, Karl R Clauser, Michael A Gillette, and Steven A Carr. Quantitative, multiplexed workflow for deep analysis of human blood plasma and biomarker discovery by mass spectrometry. Nature Protocols, 12(8):1683, 2017.

[3] Luke F Vistain and Savaş Tay. Single-cell proteomics. Trends in Biochemical Sciences, in press, doi: 10.1016/j.tibs.2021.01.013, 2021.

[4] Harrison Specht, Edward Emmott, Aleksandra A Petelski, R Gray Huffman, David H Perlman, Marco Serra, Peter Kharchenko, Antonius Koller, and Nikolai Slavov. Single-cell proteomic and transcriptomic analysis of macrophage heterogeneity using SCoPE2. Genome Biology, 22(1):1–27, 2021.

[5] Pier Giorgio Righetti and Egisto Boschetti. Introducing Low-Abundance Species in Proteome Analysis. In Low-Abundance Proteome Discovery, pages 1–11. Elsevier, jan 2013.

[6] Yao Yao, Margreet Docter, Jetty Van Ginkel, Dick de Ridder, and Chirlmin Joo. Single-molecule protein sequencing through fingerprinting: computational assessment. Physical Biology, 12(5):055003, 2015.

[7] Jagannath Swaminathan, Alexander A Boulgakov, Erik T Hernandez, Angela M Bardo, James L Bachman, Joseph Marotta, Amber M Johnson, Eric V Anslyn, and Edward M Marcotte. Highly parallel single-molecule identification of proteins in zeptomole-scale mixtures. Nature Biotechnology, 36(11):1076–1082, 2018.

[8] Shilo Ohayon, Arik Girsault, Maisa Nasser, Shai Shen-Orr, and Amit Meller. Simulation of single-protein nanopore sensing shows feasibility for whole-proteome identification. PLoS Computational Biology, 15(5):e1007067, 2019.

[9] Nicolas Abello, Huib AM Kerstjens, Dirkje S Postma, and Rainer Bischoff. Selective acylation of primary amines in peptides and proteins. Journal of Proteome Research, 6(12):4770–4776, 2007.

[10] Jagpreet S Nanda and Jon R Lorsch. Labeling of a protein with fluorophores using maleimide derivitization. Methods in Enzymology, 536:79–86, 2014.

[11] Hadjer Ouldali, Kumar Sarthak, Tobias Ensslen, Fabien Piguet, Philippe Manivet, Juan Pelta, Jan C Behrends, Aleksei Aksimentiev, and Abdelghani Oukhaled. Electrical recognition of the twenty proteinogenic amino acids using an aerolysin nanopore. Nature Biotechnology, 38(2):176–181, 2020.

[12] Gang Huang, Kherim Willems, Misha Soskine, Carsten Wloka, and Giovanni Maglia. Electro-osmotic capture and ionic discrimination of peptide and protein biomarkers with FraC nanopores. Nature Communications, 8(1):1–11, 2017.

[13] Shengli Zhang, Gang Huang, Roderick Versloot, Bart Marlon Herwig, Paulo Cesar Telles de Souza, Siewert-Jan Marrink, and Giovanni Maglia. Bottom-up fabrication of a multi-component nanopore sensor that unfolds, processes and recognizes single proteins. bioRxiv, doi: 10.1101/2020.12.04.411884, 2020.

[14] Gang Huang, Arnout Voet, and Giovanni Maglia. Frac nanopores with adjustable diameter identify the mass of opposite-charge peptides with 44 dalton resolution. Nature Communications, 10(1):1–10, 2019.

[15] Laura Restrepo-Pérez, Chirlmin Joo, and Cees Dekker. Paving the way to single-molecule protein sequencing. Nature Nanotechnology, 13(9):786–796, 2018.

[16] Ruedi Aebersold, Jeffrey N Agar, I Jonathan Amster, Mark S Baker, Carolyn R Bertozzi, Emily S Boja, Catherine E Costello, Benjamin F Cravatt, Catherine Fenselau, Benjamin A Garcia, et al. How many human proteoforms are there? Nature Chemical Biology, 14(3):206, 2018.

[17] Erik C Yusko, Brandon R Bruhn, Olivia M Eggenberger, Jared Houghtaling, Ryan C Rollings, Nathan C Walsh, Santoshi Nandivada, Mariya Pindrus, Adam R Hall, David Sept, et al. Real-time shape approximation and fingerprinting of single proteins using a nanopore. Nature Nanotechnology, 12(4):360–367, 2017.

[18] Fabien Piguet, Hadjer Ouldali, Manuela Pastoriza-Gallego, Philippe Manivet, Juan Pelta, and Abdelghani Oukhaled. Identification of single amino acid differences in uniformly charged homopolymeric peptides with aerolysin nanopore. Nature Communications, 9(1):1–13, 2018.

